# Secretin modulates appetite via brown adipose tissue - brain axis

**DOI:** 10.1101/2022.05.26.493657

**Authors:** Lihua Sun, Sanna Laurila, Minna Lahesmaa, Eleni Rebelos, Kirsi A. Virtanen, Katharina Schnabl, Martin Klingenspor, Lauri Nummenmaa, Pirjo Nuutila

## Abstract

Secretin activates brown adipose tissue (BAT) and induces satiation in both mice and humans. However, the exact brain mechanism of this satiety inducing, secretin-mediated gut-BAT-brain axis is unknown. In this placebo-controlled, single-blinded neuroimaging study, firstly using [^18^F]FDG-PET measures (n = 15), we established that secretin modulated brain glucose consumption through the BAT-brain axis. Predominantly, we found that BAT and caudate glucose uptake levels were negatively correlated (r = −0.54, p = 0.037) during secretin but not placebo condition. Then, using functional magnetic resonance imaging (fMRI; n = 14), we found that secretin down-regulated the brain response to appetizing food images and improved inhibitory control. Finally, in a PET-fMRI fusion analysis (n = 10), we disclosed the patterned correspondence between caudate glucose uptake and neuroactivity to reward and inhibition, showing that the secretin-induced neurometabolic coupling pattern promoted satiation. These findings suggest that secretin modulates the BAT-brain metabolic crosstalk and subsequent neurometabolic coupling to induce satiation, bearing potential clinical benefits for treating eating disorders.

**Significance of the study:** Secretin activates brown adipose tissue and induces satiation, but the underlying brain mechanisms are still unclear. This placebo-controlled PET-fMRI study uses brain metabolic and BOLD measures to dissect the modulatory effects of secretin on brain functions associative to satiation. Findings show that secretin i) modulates caudate glucose metabolism via the BAT-brain axis, ii) enhances BOLD response in inhibitory control, and iii) reduces reward-related BOLD response. Further evidence shows that these measured effects are tightly linked via the secretin-mediated brain neurometabolic coupling. This study significantly advances our knowledge on how secretin leads to satiation and highlights the potential role of secretin in treating eating disorders and obesity.

## Introduction

Secretin is the first hormone ever discovered, and its best-known effect is the induction of pancreatic exocrine secretion (Bayliss and Starling, 1902). It is secreted while having a meal and has recently been found to induce satiation in both mice and humans (Li et al., 2018; Laurila et al., 2021). It is suggested that secretin-induced satiation may occur by activation of POMCergic neurons in the medio-basal hypothalamus, thus targeting homeostatic circuits involved in the regulation of food intake (Cheng et al., 2011; Jeong et al., 2018; Li et al., 2018). According to the thermoregulatory feeding theory (Himms-Hagen, 1995), thermogenesis in the brown adipose tissue (BAT) is detected by hypothalamic thermo-sensors modulating central regulation of food intake (Li et al., 2018). Indeed, satiation induced by secretin depends on the activation of BAT thermogenesis where secretin binds to its widely expressed receptors in BAT to activate thermogenesis (Li et al., 2018). This delivers a satiation-stimulating signal to the brain, which may be conveyed by heat-mediated BAT-brain metabolic crosstalk, as suggested previously, but the involvement of endocrine and neuronal communication cannot be excluded (Li et al., 2018). The signalling mode(s) between BAT and brain as well as the targeted brain regions remain to be elucidated in more detail (Schnabl et al., 2021).

As the sympathetic nerves controlling BAT thermogenesis harbour afferent sensory fibers projecting to the brainstem, the midbrain, and the forebrain (Bartness et al., 2010). Secretin-induced BAT activation may trigger afferent neuronal communication with multiple brain regions. We have previously shown in humans, that secretin quenches brain reward-related BOLD responses to appetizing food and increases BAT glucose consumption (Laurila et al., 2021). The observed neuroanatomical localization of BOLD responses further suggests modulation of the limbic reward system and cognitive control in secretin-induced satiation. In the meanwhile, altered brain glucose metabolism, such as the increased caudate glucose uptake, has been linked with binge eating behavior (Nummenmaa et al., 2012; Tuulari et al., 2013). Hence, it is justified to probe whether secretin, via the BAT-brain axis, further affects the brain glucose metabolism and consequently the neurometabolic coupling with cerebral BOLD signals associative to satiation.

Satiation is linked with altered patterns of neural activations in the cortex (Thomas et al., 2015; Laurila et al., 2021). Our previous study especially shows that secretin-mediated satiation delivers a suppressive effect on brain BOLD responses to appetizing food cues. Conversely, elevated neural activation to food cues has been shown to be increased in obese subjects (Nummenmaa et al., 2012), suggesting a shared neural basis for secretin-induced satiation and trait-level eating behaviour. While we have shown that secretin downregulates reward-related neural BOLD responses, it is still unclear whether this effect is simply due to reduced sensitivity to visual food stimuli, or whether it is also accompanied by increased cognitive control that suppresses a motivation to eat. Aberrant brain activity in response inhibition has been closely linked with trait-level binge eating (Lock et al., 2011; Hege et al., 2015). Therefore, it is possible that secretin leads to satiation also via modulating brain inhibitory control in healthy subjects. However, this remains to be explored.

Brain BOLD activity is tightly linked with regional brain glucose metabolism at both resting state (Tomasi et al., 2013; Aiello et al., 2015) and during tasks, yet with markedly different patterns of association (Stiernman et al., 2021). More specifically, while a largely positive regional association was found in resting states, neurometabolic coupling during cognitive tasks suggests dissociations between the two dimensions of measures. Neuroimaging of satiation and fasting states suggests varying cortical BOLD responses (Thomas et al., 2015; Laurila et al., 2021), and therefore neurometabolic coupling between cortical BOLD and metabolic supply to the central reward hub may naturally possess varying patterns. However, to our knowledge, no previous studies have investigated these aspects. Understanding the patterns of neurometabolic coupling linked to satiation will expand the current knowledge on the brain mechanisms of eating behaviour. The current study also aims to provide novel information on these issues that are specific to secretin-induced satiation.

In the current exploratory study, we specifically investigated the central mechanism of secretin mediated satiation using our placebo controlled GUTBAT Trial dataset (Laurila et al., 2021). We first investigated whether the secretin-mediated BAT glucose uptake was correlated with corresponding brain glucose consumption, determining whether secretin modulates brain glucose metabolism through the BAT-brain axis. We then studied whether secretin modulates brain inhibitory control by measuring cortical BOLD signals during a response inhibition task. Finally, we examined whether the secretin-sensitive metabolic supply is directly linked with the secretin-modulated brain inhibitory function and food reward responses, using PET-fMRI fusion analysis. We hypothesize that secretin modulates glucose metabolism in central reward hubs through the BAT-brain axis. We further hypothesize that the secretin-sensitive metabolic supply, via gearing neurometabolic coupling, affects cognitive control and reward processing leading to satiation.

## Methods

### Study design

We investigated the effects of intravenous secretin infusions on BAT and brain metabolism (measured with [^18^F]FDG PET), as well as brain BOLD responses to food-reward images and inhibitory control (measured with fMRI). This randomized crossover study was placebo-controlled, and the participants were blinded to the intervention. All scans were performed in the morning after an overnight fast and at room temperature. Repeated PET measures during placebo and secretin conditions had an interval of 2-30 days (median 15 days) and fMRI 7-30 days (median 14.5 days). The study protocol was reviewed and approved by the Ethics Committee of the Hospital District of Southwest Finland. The PET/CT trial was prospectively registered in the EudraCT registry 2.6.2016 (EudraCT Number: 2016-002373-35) and a major amendment which included the fMRI study was registered prospectively in Clinical Trials registry 25.9.2017 (Clinical Trials no. NCT03290846). The primary and secondary endpoints of the trial have been reported previously, where more details of the study design are available (Laurila et al., 2021).

### Subjects

Fifteen healthy male participants [mean (s.d.) Age 41.6 ±12.1 years, BMI 23.6 ±1.9 kg/m^-2^] took part in the [^18^F]FDG PET brain imaging study. In parallel, fourteen healthy male participants (Age 34.4 ±14.6 years, BMI 23.3 ±1.8 kg/m^-2^) joined the fMRI study. Among these participants, a total of 10 males were studied with both [^18^F]FDG PET and fMRI. The experiment was conducted according to the declaration of Helsinki and all participants provided written informed consent for participating in the study (Clinical Trials no NTC03290846).

### BAT PET data acquisition and processing

The [^18^F]FDG-PET scans were conducted using GE Discovery (GE DiscoveryTM ST System, General Electric Systems) as described previously (Laurila et al., 2021). Firstly, a CT scan of the neck was performed for anatomic localization. Next, 150 MBq of [^18^F]FDG was administered for measuring GU (Orava et al., 2011) and a second two minute infusion of placebo or secretin was initiated. Dynamic 40 minutes scanning was started simultaneously on the neck region (frames: 1 × 1 min, 6 × 30 sec, 1 × 1 min, 3 × 5 min and 2 × 10 min). Arterialized venous plasma radioactivity samples were collected during the scan. Radiotracer [^18^F]FDG was produced using FASTlab synthesis platform (GE Healthcare) as previously described (Hamacher et al., 1986).

Image analysis was conducted with Carimas 2.8 software (Turku PET Centre, Turku, Finland). Regions of interest (ROI) were manually outlined in the fusion images, composed of the dynamic [^18^F]FDG PET image and the corresponding CT image. To analyse BAT GU, ROIs were drawn on the supraclavicular fat depots including only voxels with CT Hounsfield Units (HU) within the adipose tissue range (−50 to −250 HU) (Din et al., 2017). For tissue glucose uptake calculations, time activity curves (TAC) were generated for the ROIs. Regional TAC data was analysed by taking into account the radioactivity in arterialized plasma using the Patlak model (Patlak and Blasberg, 1985). A lumped constant value of 1.14 was used for adipose tissue (Virtanen et al., 2001).

### Brain PET data acquisition and processing

Dynamic 15 minute brain PET scans (frames: 3 × 5 min) were started 70 minutes after [^18^F]FDG injection. Head movement was prevented by strapping to the scan table. Computed tomography scans were obtained prior to the PET scans for attenuation correction and the T1-weighted MR images (TR 8.1 ms, TE 3.7 ms, flip angle 7°, 256 × 56 × 176 mm^3^ FOV, 1 × 1 × 1 mm^3^ voxel size) were taken using the 3-Tesla Philips Ingenuity PET/MR scanner for anatomical normalization and reference. PET data were preprocessed by the automatic pipeline Magia (Karjalainen et al., 2020). PET brain images were motion-corrected and then coregistered to the corresponding structural MR images. Brain glucose uptake was estimated using fractional uptake rate (Thie, 1995).

### Inhibitory control task

Participants were instructed to press a button using the left hand when it was a go signal or to withhold from pressing the button when it was a nogo signal (**Fig 1B**), while brain haemodynamic responses were measured. This task was performed immediately after the anticipatory food-reward task, see the supplementary data and our previous study (Laurila et al., 2021). Small dots were presented one-by-one in the middle of the computer screen, with an interval of 0.8 seconds. The dots could be either grey (go signal, 70% of all cases), green (nogo or go signal, 15% of all cases) or blue nogo or go signal, 15% of all cases). The grey dots were always “go” signal prompting the subject to press the button. The blue and green dots were randomly assigned to either rare “go” or “no go” signal for each participant. The statistical contrast for inhibitory activation by either blue or green dots occurred with equal probability. Stimulus delivery was controlled by the Presentation software (Neurobehavioral System, Inc., Berkeley, CA, USA).

**Figure 1.**
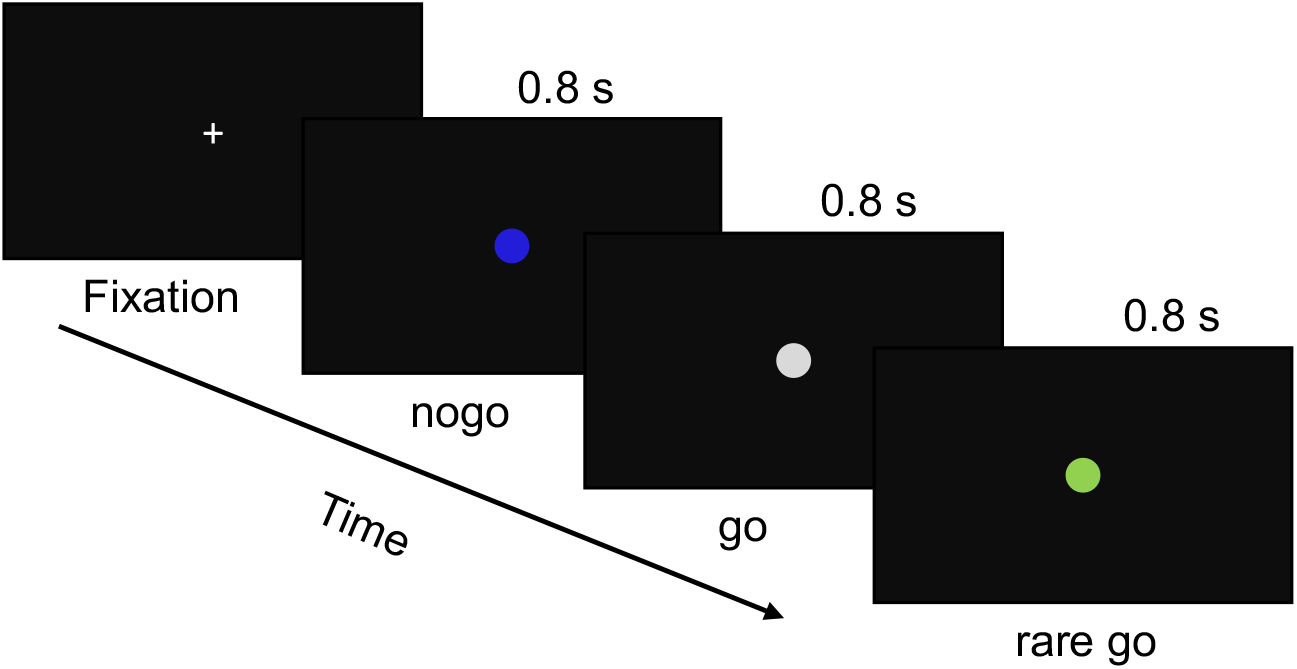
fMRI paradigm for the response inhibition task. Subjects were instructed to press the button on the go and rare go signals, and refrain from pressing the button on the nogo signals.

### fMRI data acquisition and processing

The Phillips Ingenuity TF PET/MR 3T whole-body scanner was used to collect MRI data. Structural brain images with resolution of 1 mm^3^ were acquired using a T1-weighted sequence (TR 9.8 ms, TE 4.6 ms, flip angle 7°, 250 mm FOV, 256 × 256 reconstruction matrix). Functional MRI data were acquired using a T2*-weighted echo-planar imaging sequence (TR = 2600 ms, TE = 30 ms, 75° flip angle, 240 mm FOV, 80 × 80 reconstruction matrix, 62.5 kHz bandwidth, 3.0 mm slice thickness, 45 interleaved slices acquired in ascending order without gaps). A total of 145 functional volumes were acquired during the inhibitory control task.

MRI data were processed using the fMRIPrep 1.3.0.2 (Esteban et al., 2019). Structural T1 images were processed following steps: correction for intensity non-uniformity, skull-stripping, brain surface reconstruction, spatial normalization to the ICBM 152 Nonlinear Asymmetrical template version 2009c (Fonov et al., 2009) using nonlinear registration with antsRegistration (ANTs 2.2.0) and brain tissue segmentation. Functional MRI data were processed in following steps: co-registration to the T1 reference image, slice-time correction, spatial smoothing with a 6mm Gaussian kernel, automatic removal of motion artifacts using ICA-AROMA (Pruim et al., 2015) and resampling to the MNI152NLin2009cAsym standard space. Quality of images was inspected visually for the whole-brain field of view coverage, proper alignment to the anatomical images, and signal artifacts, and inspected also via the visual reports of fMRIPrep. We set to exclude images having large movement artefacts with more than 25% of the volumes exceeding 0.5-mm framewise displacement (Parkes et al., 2018)., and accordingly all functional data were included in the current study.

### Statistical analysis

#### PET data

The full-volume brain PET data were analysed using SPM12 (Wellcome Trust Center for Imaging, London, UK; http://www.fil.ion.ucl.ac.uk/spm). Firstly, paired-T test (secretin vs. placebo conditions) was used to examine the effect of secretin on brain GU. Secondly, paired-T test was done while controlling for covariates of BAT GU, where the between-condition contrast essentially indicated an interaction effect between Condition (placebo vs. secretin) and the covariates. This was used to evaluate the modulatory effect of secretin on the BAT-Brain metabolic crosstalk regarding GU. Considering the dependency between Condition and BAT GU, statistical threshold was set at p < 0.001 with FDR cluster level correction to minimize potential false positive findings. Next, GLMs using BAT GU as regressor, separately for placebo and secretin conditions, were constructed to examine the correlation between BAT GU and brain GU. Statistical threshold was set at p < 0.05 with FDR cluster level correction. In addition, correlation between BAT GU and caudate GU at caudate were done using Kendall correlation test using R statistical software (version 3.6.3). Caudate GU was estimated using MarsBarR toolbox (Brett et al., 2002) based on the ROI defined by the AAL atlas (Tzourio-Mazoyer et al., 2002).

#### fMRI data analysis

Reaction times for the go trials (either including or excluding rare go trials) were analysed and those below 100 ms or over 800 ms were excluded. Accuracy rates were estimated as the percentage of “no response” in all nogo trials. RTs and accuracy rates were analysed separately using mixed effect linear model with Condition as the fixed factor and Subject as random factor. All analysis were done using R statistical software.

The full-volume fMRI data were analysed in SPM12. The whole-brain random effects model was applied using a two-stage process with separate first and second levels. For each subject, first-level GLM was used to predict regional effects of task parameters (nogo vs. go signals; go signals including both “go” and “rare go” signals) on BOLD indices of activation and data from both conditions (secretin vs. placebo) were fitted into the same model. Statistical threshold was set at p < 0.05 with FDR correction at cluster level.

ROIs including the insula and motor area (including the presupplementary motor area, pre-and post-central cortex) were selected based on previously validated involvement in response inhibition, for example (Sebastian et al., 2013). ROI values were estimated using MarsBarR toolbox based on the ROI defined by the AAL atlas. ROI data were estimated and fitted to linear regression models using Condition and Trial type (go vs. nogo) as factors.

#### PET-fMRI fusion analysis

The fusion analysis was done in SPM12. GLMs using caudate GU as regressors, separately for placebo and secretin conditions, were constructed to examine the correlation between caudate GU and cerebral BOLD signal during either inhibitory or food-reward responses. Statistical threshold was set at p < 0.05, FDR corrected at cluster level.

## Results

### Effect of secretin on BAT and Brain GU metabolic crosstalk

Full-volume analysis of brain GU levels revealed no statistically significant difference between the placebo and secretin conditions. In contrast, there was a significant difference between conditions while controlling for BAT GU, suggesting an interaction effect between Condition (placebo vs. secretin) and BAT GU levels on GU levels in the brain (**Fig. 2A**). This may highlight a widely spreading interference of secretin on the BAT and brain metabolic crosstalk.

**Figure 2.**
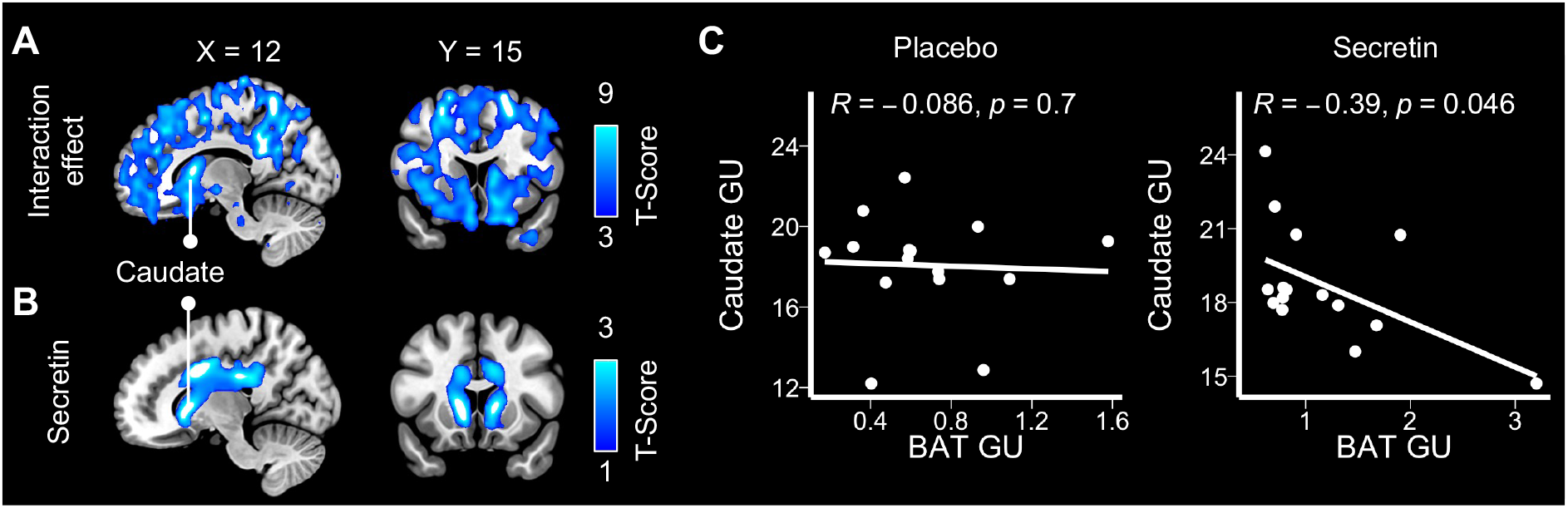
Secretin modulated the BAT-brain metabolic crosstalk (n = 15). **A)** Brain regions where secretin impacts BAT GU and brain GU associations. This was illustrated by an interaction effect between Condition (secretin vs. placebo) and BAT GU on the brain GU. Data were FDR-thresholded at p < 0.001. **B)** BAT GU was a significant predictor for GU in the brain caudate and cingulate cortex in the secretin condition, but not in any brain area in the placebo condition. Data were FDR-thresholded at p < 0.05. **C)** ROI analysis showed that GU at caudate and BAT were negatively correlated during the secretin condition but not in the placebo condition. BAT = brown adipose tissue, GU = glucose uptake.

Further analysis revealed that BAT GU was a significant predictor for GU in the caudate and cingulate area in the secretin condition (**Fig. 2B**), but not a significant predictor for GU in any brain area in the placebo condition. Further, ROI level analysis also revealed that caudate GU was negatively correlated with BAT GU in the secretin but not placebo condition (**Fig 2C**).

### Effect of secretin on inhibitory neural responses

Compared to placebo condition, secretin condition was associated with faster reaction speed in Go trials while including both “go” or “rare go” trials as go trials in the analysis (β = −0.03, 95% CI [−0.05, −0.01]). When the “rare go” trails were excluded, secretin was similarly associated with faster reaction speed (β = −0.02, 95% CI [−0.04, −0.0004]). There was no secretin-dependent effect on accuracy of responses (β = −0.06, 95% CI [−0.13, 0.02]) between the two conditions.

Brain BOLD signals in nogo versus go trials were associated with elevated activity in insula, supplementary motor area, precentral gyrus, postcentral gyrus, anterior and middle cingulate cortex, precuneus and thalamus in the placebo conditions (**Fig. 3A**). In the secretin condition, elevated activity in these brain regions were enhanced (**Fig. 3B**), as was also confirmed by an interaction effect between Condition and Type of trials (nogo vs. go; **Fig. 3C**). The ROI analysis yielded corroborating findings (**Fig. 3D**). While no significant interaction effects were found between Condition and Type of trials, in the secretin condition, nogo trials were associated with significantly increased BOLD signal in sensory and motor area (beta = 0.76, 95% CI [0.08, 1.45], p = 0.03) and insula (beta = 0.83, 95% CI [0.13, 1.52], p = 0.02). No similar effects were found in the placebo condition.

**Figure 3.**
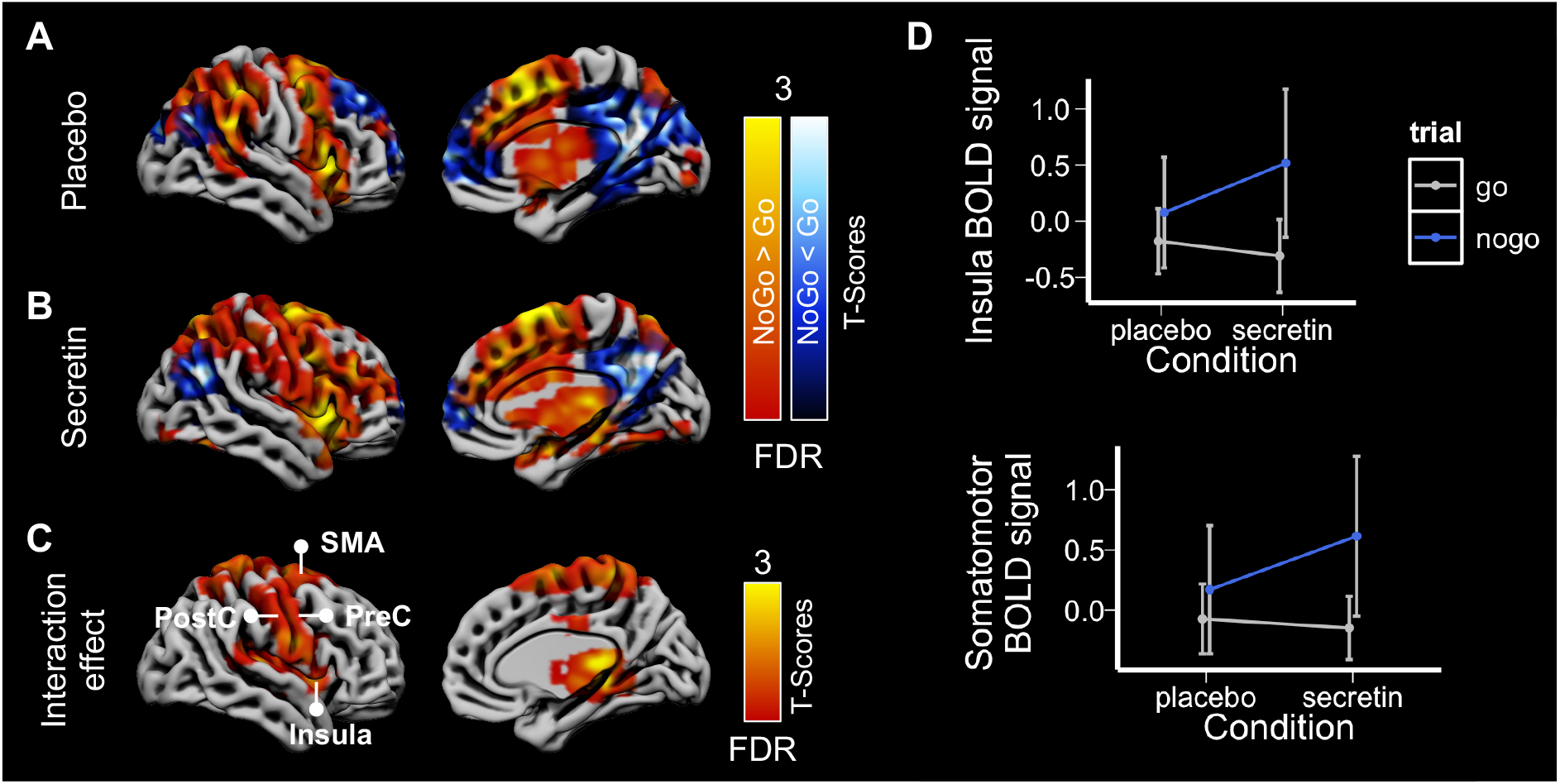
Secretin modulated the inhibition related neural activity (n = 14). Contrast images demonstrated the increased and decreased neural activity during inhibition in **A)** the placebo condition and **B)** the secretin condition. **C)** Interaction contrast between Type of trials (nogo vs. go trials) and Condition (placebo vs. secretion) showed the modulatory effect of secretin on inhibition-related neural activity. Full-volume analysis data were FDR-thresholded at p < 0.05 and right hemispheres are presented for illustration. **D)** ROI analysis showed an increased neural activity during inhibition in the motor area (comprising the supplementary motor area, precentral and postcentral gyri) and insula in the secretin condition. Error bars represent 95% Confidence Interval. BOLD = Blood oxygen level–dependent, SMA = supplementary motor area, PostC = postcentral cortex, PreC = precentral cortex.

Meanwhile, dampened activity was found in lateral and medial prefrontal cortex, and posterior cingulate area in both conditions (**Fig. 3A&B**)

Our findings so far have revealed that secretin i) stimulates metabolic crosstalk between BAT and caudate, ii) increases brain BOLD signals in inhibitory control, and ii) reduces BOLD response to appetizing food pictures (supplementary **Figure 5-1**) (Laurila et al., 2021). Next, we investigated whether these effects are tightly linked, by studying possible secretin-dependent brain neurometabolic coupling. In the following session, we reported i) whether secretin modulated correspondence between caudate GU and inhibitory BOLD responses and then ii) whether secretin modulated correspondence between caudate GU and reward-related BOLD responses.

### Correspondence between caudate glucose uptake and inhibitory responses

First-level BOLD contrast images (nogo vs. go) were analysed using caudate GU as predictor. In the placebo condition, caudate GU levels were associated with increased neural activity in the lateral prefrontal cortex, anterior- and mid-cingulate, and insula; also, caudate GU levels were associated with reduced activity in the parietal and occipital areas (**Fig. 4A**). In the secretin condition, caudate GU levels were associated with globally reduced BOLD activities (**Fig. 4B**).

**Figure 4.**
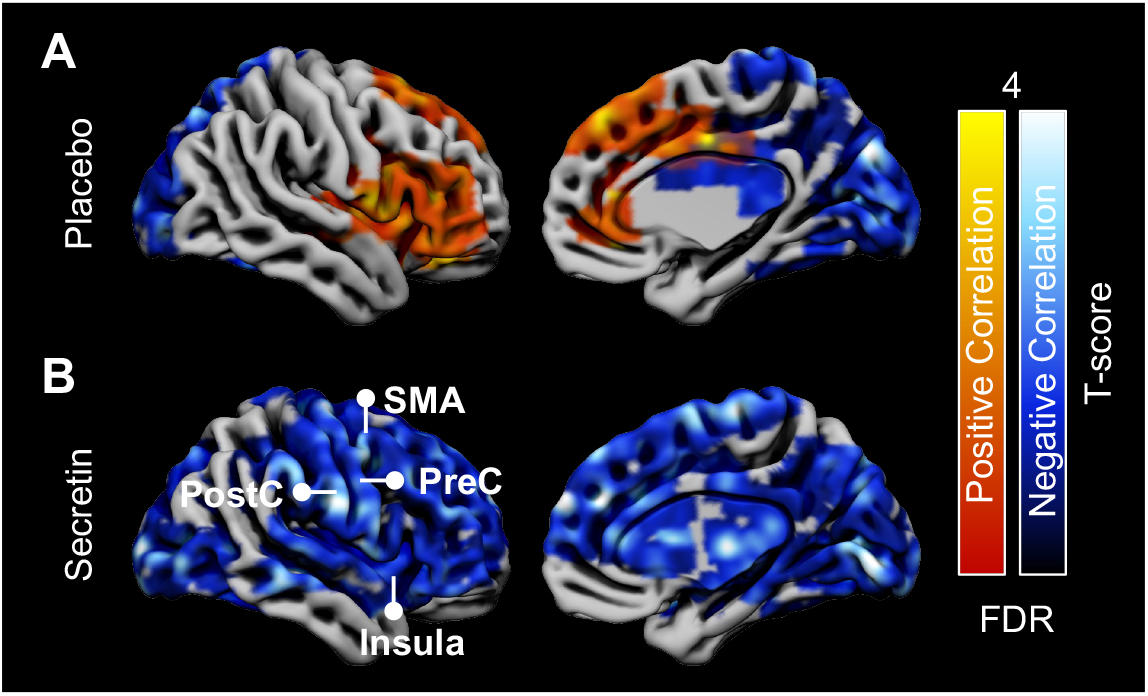
Secretin modulated the correspondence between caudate GU and cerebral BOLD during inhibition (n = 10). **A**) In the placebo condition there were both positive and negative associations between caudate GU and cerebral BOLD during inhibition. **B**) In the secretin condition, there was only a negative association. Data were FDR-thresholded at p < 0.05.

### Correspondence between caudate glucose uptake and neural activity during reward response

First-level BOLD contrast images (appetizing vs. bland food) were analysed using caudate GU as predictor. In the placebo condition caudate GU levels were associated with increased neural activity in the lateral prefrontal cortex, insula and occipital cortex; also, they were associated with reduced activity in the post-central and parietal areas (**Fig. 5A**). In the secretin condition, caudate GU levels were only associated with increased BOLD activity, but this effect was restricted to the para-central area (**Fig. 5B**).

**Figure 5.**
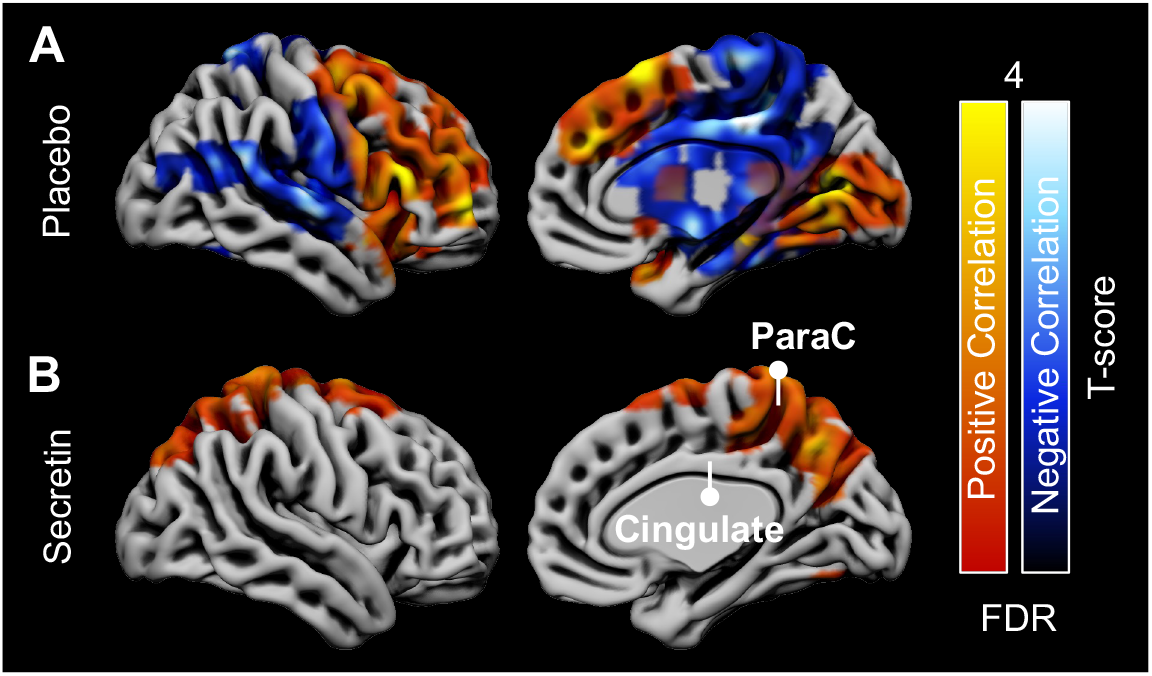
Secretin modulated the correspondence between caudate GU and cerebral BOLD signals during food-reward responses (n = 10). **A**) In the placebo condition, there were both positive and negative associations between caudate GU and cerebral BOLD signals in reward response. **B**) In the secretin condition, there was only a positive association. Data were FDR-thresholded atp < 0.05. ParaC = Paracentral gyrus.

## Discussion

The current study reveals *in vivo* that secretin induces metabolic BAT-brain crosstalk measured by [^18^F]FDG PET/CT. Using functional measures of neuroactivity, our study also shows that secretin directly shapes central food-reward responses and cognitive control. Our data display secretin-specific patterns of neurometabolic coupling during both reward responses and inhibition. Taken together, we propose that secretin modulates BAT-brain metabolic crosstalk and subsequently shapes neurometabolic coupling to generate the cognitive concepts of satiation. This study uncovers a potential brain mechanism for secretin-induced satiation, bearing prospective clinical significance in dealing with eating disorders.

Our previous studies have highlighted that BAT has a dual role in maintaining energy homeostasis (Li et al., 2018; Laurila et al., 2021). It is not merely a heater organ that increases energy expenditure, but also regulates energy intake. In both mice and men, feeding activates BAT, but the concept of thermoregulatory feeding has not been investigated in detail, especially in humans. Our previous study has introduced a neurobiological concept for heat induced satiation in mice (Li et al., 2018). Since detailed methods used in mouse models are not fully translatable to humans, the question whether BAT conveys its satiation effect to the central nervous system through heat or other means, has remained unresolved. Here, our results showed that secretin induced crosstalk between BAT and brain, which further supported the significance of a gut-BAT-brain axis in controlling eating behaviour.

The role of the caudate in reward-oriented action has been well established (Hikosaka et al., 1989; Tricomi et al., 2004). Prior studies have also shown that caudate glucose uptake is up-regulated in obese subjects, evidenced by [^18^F]FDG-PET measures during hyper-insulinemic clamp (Nummenmaa et al., 2012; Tuulari et al., 2013). Bariatric surgery attenuates this effect (Tuulari et al., 2013), which further highlights its potential role in motivation of eating. Interestingly, our findings show a link between secretin-stimulated BAT glucose uptake and the downregulation of caudate metabolism. Secretin induced higher BAT glucose uptake, which subsequently led to lower caudate glucose uptake. Lower caudate glucose supply was also associated with lower BOLD response to rewarding food images (i.e., enhanced positive correlation) and higher BOLD response in inhibition (i.e., enhanced negative correlation). In contrast, this type of correspondence was not active during the placebo condition, probably due to a baseline stochastic state of glucose uptake in the brain, where even obese subjects have previously shown no difference with normal weighted controls (Nummenmaa et al., 2012; Tuulari et al., 2013).

Secretin’s suppressive effect on acute brain responses to appetizing food images and enhancement of cognitive control further highlights its potential role in weight control. Previous studies have shown that binge eating behaviour is associated with enhanced BOLD response to rewarding food images (Nummenmaa et al., 2012) and aberrant inhibitory control (Lock et al., 2011; Hege et al., 2015). Here, secretin seems to affect both these cognitive functions, along with trait-level responses such as reduced motivation to eat (Laurila et al., 2021). Furthermore, both these cognitive responses are directly linked with the secretin-mediated change in caudate glucose uptake levels, suggesting that secretin induces satiation most probably via modulating the neurometabolic coupling between caudate glucose supply and cerebral BOLD responses.

How this metabolic crosstalk between BAT and caudate occurs remains elusive. BAT is largely innervated by the central nervous system (Bartness et al., 2010; Kuruvilla, 2019), probably including the caudate as supported by findings through the anterograde transneuronal viral tract tracing (Vaughan and Bartness, 2012). Our data showing that secretin activates BAT thermogenesis and metabolic crosstalk with the caudate suggests a potential feedback loop in the BAT-brain axis. Conversely, central control of BAT activity via neurotransmitters has been also reported. For example, release of noradrenaline activates BAT adrenergic receptors thus to stimulate biochemical reactions in mitochondria and thermogenesis (Bartness et al., 2010). Also, intravenous and intracerebroventricular administration of fentanyl enhances BAT sympathetic nerve activity and thermogenesis, suggesting a potential role of central opioid signalling in BAT thermogenesis (Cao and Morrison, 2005). However, whether these neurotransmission signalling pathways are involved in secretin-mediated BAT-caudate crosstalk, subsequently associative to satiation, remains to be explored.

## Limitations

Despite showing a significant correlation between BAT and caudate glucose uptake under secretin administration, a causal link between the effects cannot be shown with the implemented method. Due to the PET-fMRI experiment design, time-scale hierarchy between these distinct levels of functional measures remains to be verified. Furthermore, considering the complexity of PET-fMRI instrumentation, the number of studied subjects, especially for the fusion analysis, was small. All subjects were normal weight males with normal glucose tolerance. Future studies are needed to further understand whether obesity is linked with abnormal function of the gut-BAT-brain axis.

## Conclusions

The current study established a secretin-mediated BAT-caudate axis in regulating human satiation. Both metabolic and cognitive level evidence suggests that secretin may be a potential drug for the treatment of eating disorders such as obesity.

## Acknowledgements

The study was conducted within the Centre of Excellence into Cardiovascular and Metabolic Diseases supported by the Academy of Finland (grant no. 307402), University of Turku, Åbo Akademi University; and funded by Turku Collegium for Science and Medicine (L.S.), the Instrumentarium Science Foundation (grant no. 190014) (S.L.), The Paulo Foundation (S.L.), Turku University Hospital Foundation (S.L.) and The Finnish Medical Foundation (grant no. 2985) (S.L.). TUM researchers were supported by the Else Kröner Fresenius Stiftung (grant no. 2017_A108 - EKFZ (M.K.) and the German Research Foundation (grant no. DFG CRC TRR333/1 BATenergy project #450149205) (M.K.).

