## supplementary Figure 5-1 for "Secretin modulates appetite via brown adipose tissue - brain axis"

### Methods

*Anticipatory food reward task*

We used a previously established task protocol for inducing anticipatory reward, by showing the participants pictures of palatable (e.g. chocolate, pizza, cakes), and bland (e.g. lentils, cereal, eggs) food pictures (Laurila et al., 2021). This task simulates situations where appetite is triggered by anticipating the actual feeding via visual food cues, such as those in advertisements. The pictures were rated in a previous study by independent participants; the ratings showed that the appetizing foods were evaluated more pleasant than the bland foods, $t(31)=4.67$, $p < 0.001$ (Nummenmaa et al., 2012).

During the task, participants viewed alternating 16.2-s epochs with pictures of palatable or non-palatable foods. Each epoch contained nine stimuli from one category, intermixed with fixation crosses. Each food stimulus was presented on either the right or the left side of the screen, and participants were instructed to indicate its location by pressing corresponding buttons. This task was used simply to ensure that participants had to pay attention to the stimuli. Stimulus delivery was controlled by the Presentation software (Neurobehavioral System, Inc., Berkeley, CA, USA).

### Results

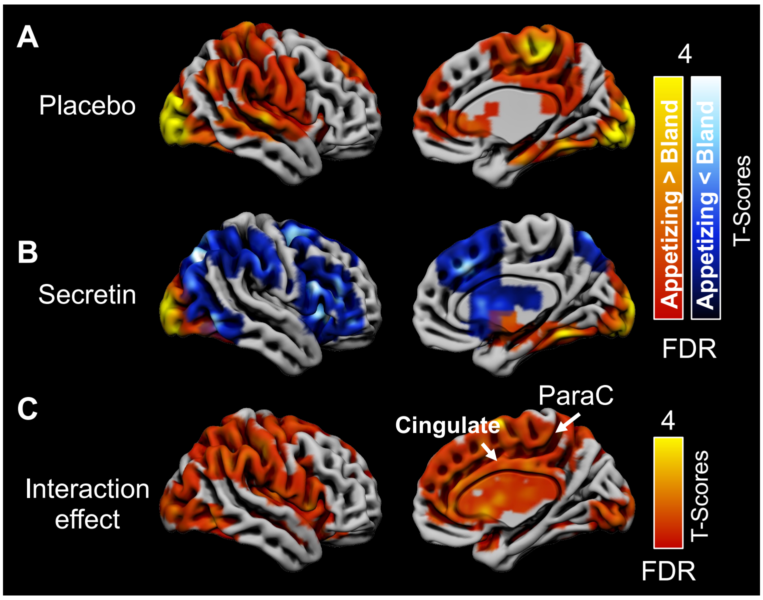


***Figure 5-1.*** *Secretin modulated the reward-related neural activity. Contrast images demonstrated the increased neural activity during inhibition in* ***A)*** *the placebo condition and both increased and dampended activity in* ***B)*** *the secretin condition.* ***C)*** *Interaction contrast between Trial types (appetizing vs. bland food images) and Condition (placebo vs. secretion) showed the modulatory effect of secretin on reward-related neural activity. Full-volume analysis data were FDR-thresholded at $p < 0.05$ and right hemispheres are presented for illustration.*
